## Supplemental Tables and Figures for "Sex-specific speed-accuracy tradeoffs shape neural processing of acoustic signals in a grasshopper"

### Supplemental Figures

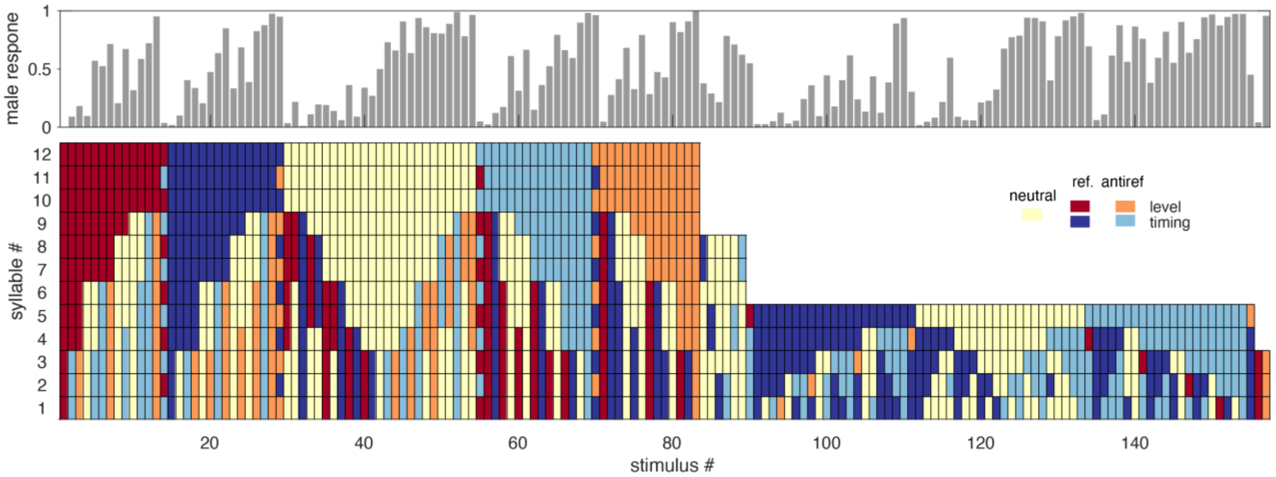


**Figure S1: Stimulus patterns (bottom, color coded, see legend) and male responses (top).**

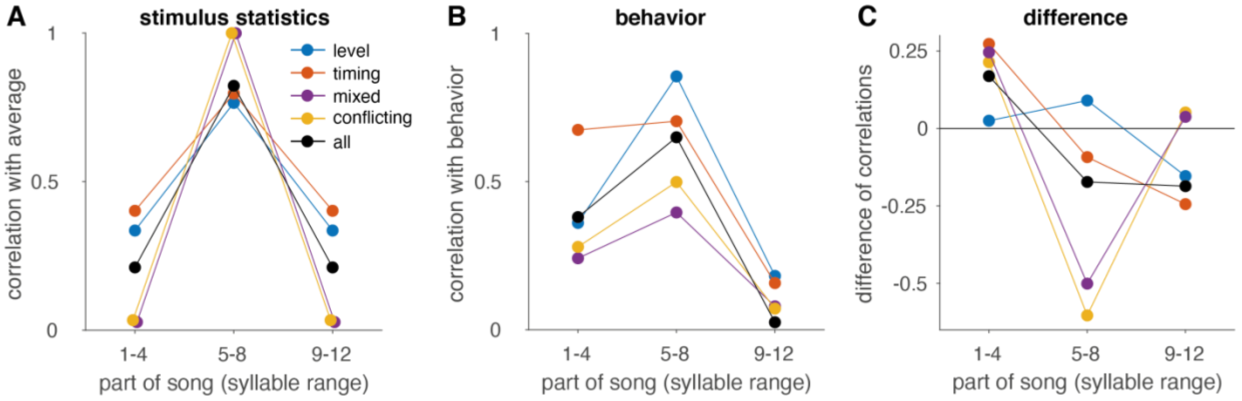


**Figure S2: Correlations in the stimulus and in the behavior.**
**A** Strength of the relationship ($r^2$) between the average directional cue from different thirds of the songs with the average directional cue over the full song (12 syllable songs only). The song's middle was more strongly correlated with the average directional cue from the stimuli compared to the beginning or the end of the song, because cue direction often changed half-way through the stimulus (see Fig. S1).
**B** Same as in A but showing the correlation between the males' turning responses and the cues from different parts of the song.
**C** Difference between the curves in B and A (see Fig. S1). The beginning of the song has a greater influence on the behavior (difference is positive) than expected from the structure of the stimuli, suggesting that males do not simply average the directional cues over the stimulus.
Calculated for different, non-exclusive sets of stimuli with 12 syllables: all (black, N=83), level only (blue, N=29), timing only (red, N=27), mixed (purple, stimuli containing timing and level cues, N=28), conflicting (orange, stimuli with cues from both sides, N=40).

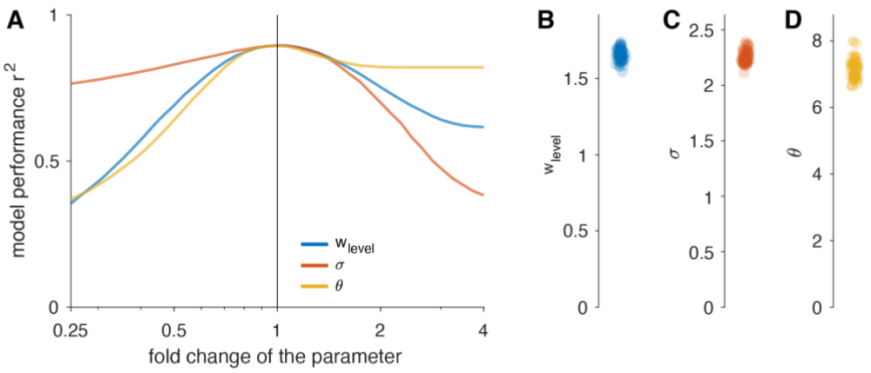


**Figure S3 – Sensitivity analysis reveals that the parameters are well determined by the data.**
**A** Performance $r^2$ of models with parameter values perturbed around the optimal ones found in Table S1. Performance decreases for changes in the parameter, indicating that the parameter values are well determined by the data. Asymmetrical effects for σ and θ arise because increasing θ does not affect the outcome for most stimuli since the sign of the integrated evidence often determines the decision even without threshold crossing. Too much noise σ likely worsens predictions for stimuli while too little noise affects model predictions more weakly.
**B, C, D** Parameter values for the weight of level cues $w_{level}$, the noise level σ, and the decision threshold θ are reproducible across cross-validation runs for the best fit model (compare Table S1).

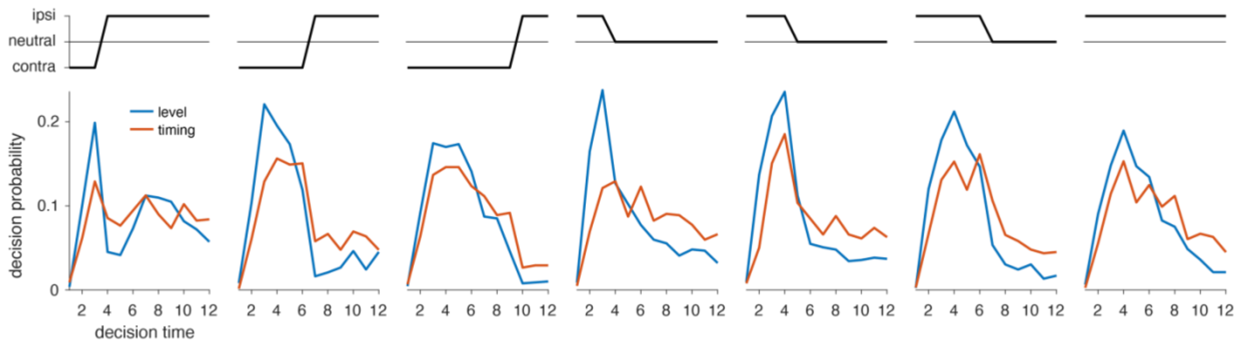


**Figure S4 – Cue sequence (top) and decision time distributions (bottom) for the stimuli in Figure 2D.**
Stimuli were chosen to have matching temporal cue sequences but different cue types (level – blue, timing – red).

| Model variant | $r^2$ | AIC | σ | θ | $w_{timing}$ | $w_{level}$ | τ | $\gamma_{urgency}$ |
|---|---|---|---|---|---|---|---|---|
| Simple averaging single cue | 0.75 | 138 | 0 | ∞ | 1 | 1 | ∞ | 0 |
| Perfect integration, single cue | 0.80 | 107 | 1.98 ± 0.05 | 5.81 ± 0.25 | 1 | 1 | ∞ | 0 |
| Perfect integration, two cues | 0.86 | 49 | 2.25 ± 0.06 | 7.14 ± 0.29 | 1 | 1.65 ± 0.04 | ∞ | 0 |
| Leaky integration, two cues | 0.86 | 53 | 1.89 ± 0.06 | 5.88 ± 0.31 | 1 | 1.67 ± 0.05 | 24.58 ± 4.81 | 0 |
| Perfect integration, two cues, urgency | 0.85 | 60 | 2.32 ± 0.12 | 7.11 ± 0.77 | 1 | 1.68 ± 0.07 | ∞ | 0.026 ± 1.01 |

**Table S1 - Model comparison and parameters.**
All values indicate the mean (± standard deviation) of parameter estimates over the 81 cross-validation runs. $r^2$ - cross-validated coefficient of determination, AIC - Akaike information criterion (smaller is better), σ - noise level, θ - decision threshold, $w_{timing}$ & $w_{level}$ - weight for timing and level cues ($w_{timing}$ fixed to 1 for all models), τ - integration time constant in units of syllables (fixed to ∞ for perfect integration), $\gamma_{urgency}$ - urgency gain (fixed to 0 for models without urgency). See also Fig. S3. Models are displayed in order of increasing complexity. The simple averaging model (first row) corresponds to a perfect, noiseless integrator without a threshold. The simplest model explaining the data best (minimal AIC score) is a drift-diffusion model with perfect integration, different weights for the two cue types, and no urgency (green shading). Simpler models (yellow shading) perform worse, in particular for stimuli with conflicting or mixed cues (Fig. 2C). More complex models (red shading) do not perform better than the best fit model.

| Property | Female | Male |
|---|---|---|
| Cost of errors | High | Low |
| Cost of Slowness | Low | High |
| Integration property | Pattern | Direction (this study) & pattern |
| Number syllables in typical input | ~30 syllables per male calling song | ~12-15 syllables per female response song |
| Time constant τ (syllables) | Very long (213 syllables) | Long (25 syllables) |
| Pos. sensory evidence w | Very weak rel. to threshold (1) | Strong rel. to threshold (1 and 1.65) |
| Neg. sensory evidence | Very strong rel. to threshold (-47) | Not tested |
| Noise σ | Very high (104) | Medium (2.25) |
| Decision threshold θ | Very high (187) | Low (7.14) |

**Table S2 – Comparison of female and male costs and integration dynamics.**
Female values from (Clemens et al., 2014).
